# The “Locus of Learning” Problem: Effects of Stimulus and Task Structure on Temporal Perceptual Learning

**DOI:** 10.1101/762633

**Authors:** Rannie Xu, Russell M. Church, Yuka Sasaki, Takeo Watanabe

## Abstract

The ability to discriminate sub-second intervals can be improved with practice, a process known as temporal perceptual learning (TPL). A central question in TPL is whether training improves the low-level sensory representation of a temporal interval or optimizes a set of task-specific response strategies. Here, we trained three groups of participants over five days on a single-interval temporal discrimination task using either fixed intervals (FI) or random intervals (RI). Before and after training, discrimination thresholds were also obtained on an untrained task. Our results revealed that only the FI group showed improvements with five days of training, but this learning did not generalize from the trained task to the untrained task in any group. These results highlight task-specificity in TPL and suggest that training-dependent improvements in timing ability might reflect an active reweighting of decision units, in addition to refinements in the sensory representation of a learned interval.

## Introduction

Temporal perceptual learning (TPL) can improve the perception and discriminability of short temporal intervals by our perceptual system. Even with a few short days of practice, performance on a variety of perceptual and motor timing tasks can be greatly enhanced (Karmarkar & Buonomano, 2003; Meegan, Aslin, & Jacobs, 2000; Westheimer, 1999; Wright et al., 1997). One of the central questions underlying these improvements concern the locus of this learning: what is being changed when we learn to time? On the one hand, TPL might reflect a refinement in the sensory representations of a trained interval (low-level hypothesis), on the other hand, TPL might involve changes in task-specific behavior which in turn lead to better performance on a trained task (high-level hypothesis). We refer to this dichotomy between low-level (i.e., learning to sense) and high-level hypotheses (i.e., learning to respond) as the “locus of learning” problem, and we aim to address this question in the present set of experiments by investigating the effects of stimulus and task structure on TPL.

Evidence of low-level changes in temporal processing following perceptual training is the finding that performance improvements are often bound to the trained interval and does not generalize to other untrained intervals (i.e., interval-specificity). In an early study by Wright and colleagues (1997), discrimination training with a 100ms auditory interval was found to significantly improve performance post-training, but the same amount of practice did not improve performance in any neighboring (e.g., 50ms, 200ms, 500ms) conditions. Interval-specificity suggests that the mechanisms underlying learning is largely temporally-specific, which hints at the existence of duration-selective tuning mechanisms in the brain (Bueti, Bahrami, & Walsh, 2008; Hayashi et al., 2015; Protopapa et al., 2018). Interval-specificity is an important characteristic of TPL and has been widely reported across the literature (Bueti et al., 2012; Hayashi et al., 2018; Nagarajan et al., 1998; van Wassenhove & Nagarajan, 2007; Wright, Wilson, & Sabin, 2010) with a few exceptions (Lapid, Ulrich, & Rammsayer, 2009b).

In contrast to the low-level hypothesis, learning can be driven alternatively by changes in the way we respond to a perceptual stimulus, optimizing the high-level decision processes specific to a trained task. In this view, the observation of learning specificity to a trained stimulus does not necessarily reflect any change in the representation of the stimulus itself (Maniglia & Seitz, 2018), suggesting that low-level learning by itself is insufficient to account for the behavioral improvements observed with practice. In a series of primate studies (Law & Gold, 2008; Law & Gold, 2009), training on a visual motion discrimination task led to changes in the lateral intraparietal area – a decision-making unit, rather than the middle temporal area, which is the primary sensory unit for representing motion. What these findings reveal about the mechanism of perceptual learning is that we must adjust for the optimal stimulus-response weights given the relevant task structure (Ahissar et al., 2009; Ahissar & Hochstein, 2004; Hochstein & Ahissar, 2002) and learning must involve changes in how sensory information is interpreted to form a behavioral choice.

The purpose of the present study is to provide evidence in support of high-level changes in temporal processing by systematically assessing the roles of stimulus and task structure on TPL. First, we ask whether learning on one timing task is transferrable to an untrained task when the intervals are identical. We hypothesized that if learning occurs at the level of interval representation (low-level), improvements on the trained interval should transfer to an unlearned task so long as the interval is identical. On the other hand if TPL relies on learning of appropriate response strategies (high-level), then we would not expect any generalization for the untrained task since there are now a different set of task structures and decision rules to be learned.

In addition, we investigated the effect of stimulus uncertainty on TPL. A recent finding on the learning of visual features (i.e., visual perceptual learning; VPL) revealed that improvements in visual discrimination relied on fixed stimulus structure during training (Adini et al., 2004; Kuai et al., 2005; Yu, Klein, & Levi, 2004). In a classic study, participants were asked to determine which stimulus pair had the highest contrast. Improvements on the contrast discrimination task was only found for the group which received practice on seven fixed base contrasts but not when they practiced the same contrast pairs in a mixed-by-trial (i.e., differing from trial-to-trial) manner. What this suggests is that the degree of stimulus uncertainty or the ability to discern statistical regularities in training input may influence the type of decision strategy that is adopted by the viewer (Adini et al., 2004). Therefore, if TPL shares similar mechanisms of learning as VPL, we would also expect roving effects to occur when we use fixed vs. random temporal stimuli during training. We predict that stochasticity (that is, higher uncertainty in a prior distribution) would impair TPL if the high-level hypothesis is true, since learning is affected by the decision strategy and sensitivity thresholds within the given task. However, if a low-level hypothesis is true, then TPL should not be affected by roving effects (or changes in stimulus structure) since learning is taking place at the sensory level for the interval.

In summary, the locus of learning problem of TPL addresses the question whether practicing a timing task primarily improves the representation of an interval (low-level hypothesis), or task-specific processing strategies (high-level hypothesis). The present study addresses this dichotomy by investigating effects of stimulus and task structure on temporal learning. Results from our experiments provide evidence in support of high-level changes in temporal discrimination performance, whereby behavioral improvements were dependent on fixed stimulus structure during training and did not generalize under a new set of task structures. Both findings are consistent with current models of perceptual learning and provide grounds for constructing a unifying framework of perceptual learning across modalities.

## Methods (Experiment 1)

### Participants

Twenty-nine right-handed young adults with normal or corrected-to-normal vision and hearing were recruited for participation in Experiment 1 (16 females; mean age: 23.8 ± 3.9 years). We justified our sample size through power calculations using the G*Power software (Mayr et al., 2007). Based on an effect size estimate of .25 and power of .80, a total of 18 subjects are needed to achieve significance. Two participants were unable to complete the experiment due to scheduling conflicts, and three more were excluded due to poor fits in their psychometric function (see *Results*) leaving a total of 24 participants in Experiment 1 (n = 12 per group). Each session was held at approximately the same time each day to avoid possible confounding effects of time of day on temporal processing (Lustig & Meck, 2001). Written consent was obtained prior to participation and approved by the institutional review board (IRB) at Brown University.

### Stimulus and Apparatus

Participants were seated in a sound-insulated room with dim lighting. All stimuli were generated and presented using MATLAB with Psychophysics Toolbox extensions, version 3.0.14 (Kleiner, Brainard, & Pelli, 2007). Visual stimuli were presented on a ViewSonic – VA2226w monitor, measuring 20 × 14 inches, with a refresh rate of 75Hz and a viewing distance of approximately 38cm. Auditory stimuli were presented at 86 dB SPL through noise-cancelling Sennheisser headphones and included a 5ms on and off ramp. All responses were collected using a standard US keyboard.

### Procedure

We used a standard pretest-training-posttest design over seven consecutive days. During pre-test and post-test days (Days 1 and 7), discrimination thresholds were estimated using an adaptive staircase procedure at four intervals (100ms, 200ms, 300ms at 1kHz and 200ms at 4kHz). The presentation order for each condition was blocked and randomized according to the Latin Square design and kept constant between pre-test and post-test sessions for each participant. The training phase (Days 2 to 6) consisted of 480 trials of practice for the 200ms at 1kHz condition using a single-interval comparison procedure, over 5 days (2400 trials in total). We justified our number of trials/training days by referring to previous studies which have demonstrated substantial learning effects using ~2500 trials (Meegan et al., 2000; Van Wassenhove & Nagarajan, 2007; Bratzke et al., 2012), and with as little as a single day of training with 900 trials (Westheimer, 1999).

### Training Phase

(Day 2 through Day 6). The training phase of Experiment 1 involves 5 consecutive days of practice on a single-interval two-alternative forced-choice temporal comparison procedure similar to the one used in Karmarkar & Buonomano (2003). At the beginning of every block, participants were presented with the to-be-learned interval (200ms) for reference and instructed to categorize subsequent comparison intervals as either “longer” or “shorter” than the reference interval. For the RI group, comparison intervals are drawn from a normal distribution (μ = 200ms, σ = 2.5μ) with limits at 158ms and 242ms. For the FI group, one of eight predetermined comparison values (158, 170, 182, 194, 206, 218, 230, 242 ms) were selected at random on every trial. These values are based on a previous study (Bratzke, Seifried, & Ulrich, 2012) and symmetrically-distributed about the 200ms reference interval. Thus, the statistical parameters of the comparison values (i.e., mean and variability) are identical between the FI and RI groups. The key difference is in whether the input structure is from a discrete (FI) or continuous (RI) distribution.

### Testing Phase

(Day 1 and Day 7). During the testing phase, a 3-up-1-down adaptive staircase procedure was used to estimate the 79% correctness on individual psychometric functions (Levitt, 1971). Two empty intervals were presented on each trial: a standard (t) and a comparison (t + Δt). Listeners must indicate which interval was the standard (i.e., shorter interval) by pressing either the “J” or “K” keys using their index and middle finger. The length of the standard and comparison was identical on the first trial of every block, forcing subjects to guess. Three consecutive “correct” responses led to a decrease in Δt, and a single “incorrect” response led to an increase in Δt. Every “correct” response immediately following an “incorrect” response, and vice versa, is referred to as a *reversal*. The discrimination threshold is estimated based on the last 6 reversals of each block. As with Wright et al. (2010), threshold estimates were only calculated if there were at least 7 total reversals within a block. The step size for the first five reversals is 5% of the standard and 1% thereafter. The inter-stimulus interval was jittered about 1000ms to minimize predictability between the offset of the first interval and the onset for the second interval.

## Results (Experiment 1)

### Training Phase

To assess the degree of learning on the trained task, we fit a psychometric function on the proportion of “long” responses, *p*(Long) = #long responses/ #total responses, at each comparison interval using the Quickpsy package in R (Linares & López-Moliner, 2019). A logistic function was used for each daily session, per subject, with lapse; all other parameters of the model were allowed to vary. Based on the goodness-of-fit indicated by the R^2^ value, we excluded data from three subjects (two from the FI group, one from the RI group) whose performance were below 2.5SD of the group average.

Next, we computed an MLE of discrimination performance (i.e., the difference limen, DL) by taking the average of the interquartile range on the fitted functions (i.e., DL = (*x*.75−*x*.25)/2). In standard 2AFC tasks, the DL can be seen as a reliable measure of sensitivity in a psychological system (Lapid, Ulrich, & Rammsayer, 2008, 2009a), and represents the least amount of change in the physical stimulus necessary for psychological indifference, *p*(Long) = .5. Therefore, a threshold value of 20ms indicates that a subject is equally-likely to respond “long” or “short” when presented with a 200ms interval against a 220ms interval. TPL is then indexed by a training-related *decrease* in discrimination threshold between the first and last day of training.

We first submitted these threshold estimates to a 2 × 2 mixed model ANOVA with group (FI/RI) as a between-subjects factor and session (first/last) as a within-subject factor. Recall that we are interested in whether the stimulus structure (i.e., being presented with fixed or random comparison values) affects the amount of improvement sustained between the groups, which would manifest as a significant interaction between the two factors. The results of our ANOVA revealed a significant main effect for session, F_1,22_ = 5.71, p = .026, η^2^ = .06. but not group, F_1,22_ = 0.03, p = .86, η^2^ = .001. In addition, we observed a significant interaction between the two factors, F_1,22_ = 4.34, p = .049, η^2^ = .05 (Figure 1). Post-hoc pairwise t-tests revealed a significant training-related decrease in discrimination thresholds on the last day of training (DL ± SD = 199 ± 8.67ms) as compared to the first day (213 ± 6.33ms; t_11_ = 3.01, p = .036). To investigate the significant interaction between training and group, we calculated a change in performance between the first and last day for both groups and performed an unpaired t-test on the two groups. This analysis revealed a significant difference between the FI group, which had a greater change in performance (mean change = 7.54ms), and the RI group (mean change = 0.48ms; t_11_ = 2.35, p = .039), demonstrating TPL in the FI group, but not the RI group.

**Figure 1.**
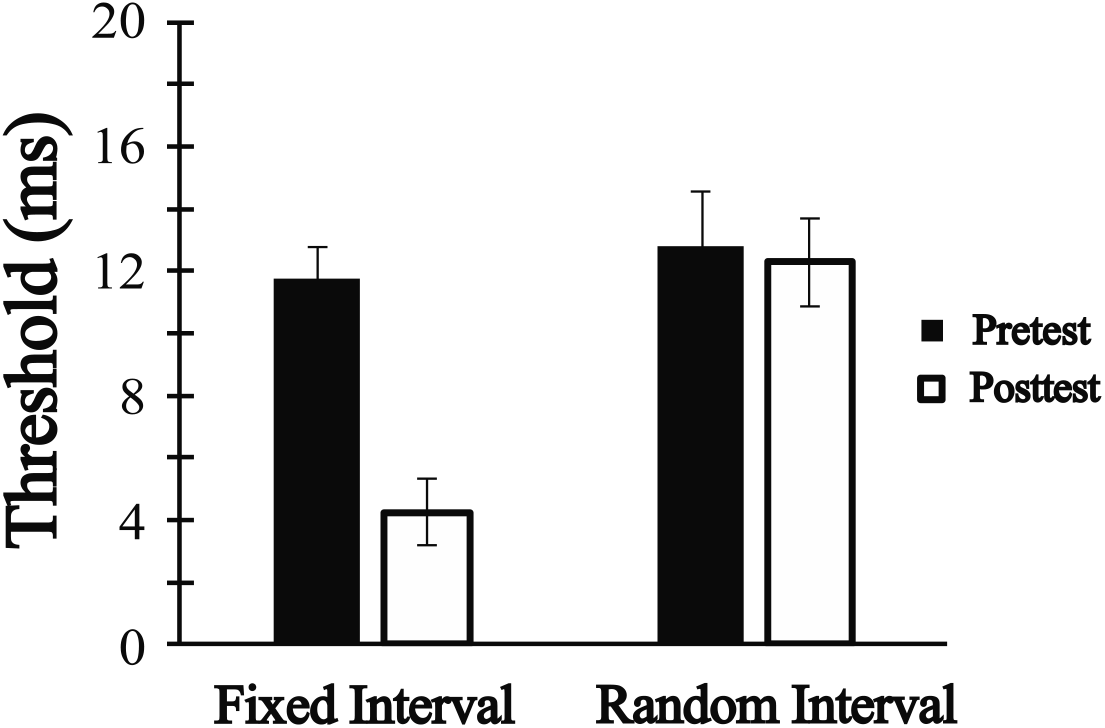
Average learning indices (change in discrimination thresholds) for the FI and RI groups before and after training. Error bars represent +/− SEM.

### Testing Phase

To obtain a comparable measure of discrimination performance in the untrained two-interval discrimination task, we also calculated a discrimination threshold for the pretest (Day 1) and posttest (Day 7) sessions based on the averaged comparison intervals in the last 6 reversals within each experimental block. Consistent with Wright et al. (2010)’s procedures, thresholds were only estimated if there were at least 7 total reversals within a block. Two estimates were obtained for each of the four experimental conditions (100ms, 200ms, 300ms at 1kHz and 200ms at 4kHz), and they were divided by their respective standard interval to obtain the Weber’s Fraction (WF) for each condition per session (Figure 2).

**Figure 2.**
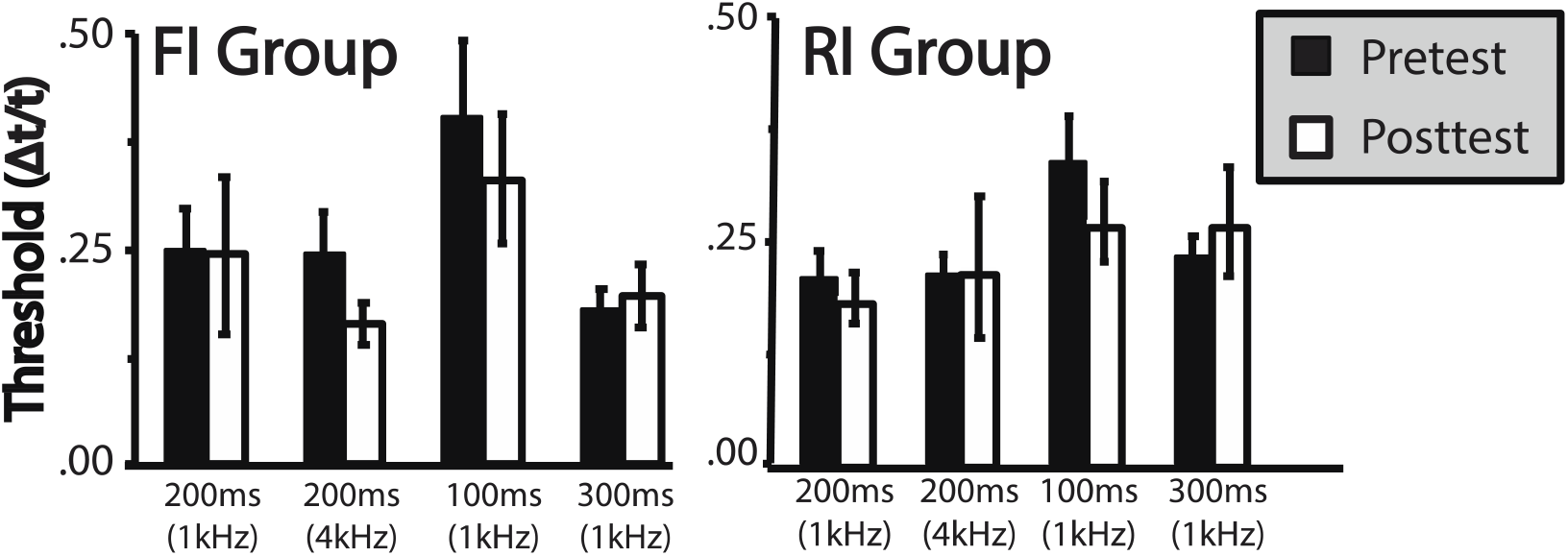
Changes in discrimination threshold between pretest and posttest sessions for the untrained task in the trained (200ms at 1kHz) and untrained conditions in Experiment 1. Error bars represent +/− SEM.

We first submitted the WFs for the FI and RI groups to a 2 group × 2 session × 4 condition mixed model ANOVA again with group (FI/RI) as a between-subjects factor, and session (pretest/posttest) and condition (100ms, 200ms, 300ms at 1kHz and 200ms at 4kHz) as within-subject factors. A significant main effect (Greenhouse-Geisser corrected) was observed for condition, F_3,66_ = 4.59, p = .034, η^2^ = .07, but not session, F_3,66_ = .88, p = .45. Post-hoc t-tests investigating the main effect of condition found that the 100ms condition had higher WFs during both the pretest (mean WF_pretest_ ± SEM = 1.29 ± .011) and posttest (mean WF_posttest_ ± SEM = 1.42 ± .026) sessions as compared to all other interval conditions. This finding is reported in a number of previous studies (e.g., Bratzke, Seifried, & Ulrich, 2012; Lapid, Ulrich, & Rammsayer, 2009b; Warm, Stutz, & Vassolo, 1975) and demonstrate the greater difficulty of discriminating shorter intervals as compared to longer ones. Importantly for the purposes of our investigation on the generalization of TPL across tasks, we did not find a significant interaction between group, condition, and session, F_3,66_ = 1.66, p = .18. This suggests that in our present study, we did not find evidence in support of a transfer of TPL between a trained and untrained task regardless of group membership (FI/RI).

## Discussion

The two main questions addressed in Experiment 1 are (1) whether TPL for a learned interval (200ms) can generalize to an unlearned task and (2) whether the stochasticity in stimulus input (i.e., fixed vs. random) during training plays a role in interval learning. First, our results demonstrate task-specificity in TPL wherein the improvements on a single-interval temporal comparison task did not transfer to a two-interval discrimination task using the same interval. Second, improvements on the single-interval comparison task was restricted to the group that received fixed input during training (FI group). We interpret the lack of learning in the RI group as a result of stochasticity in the training input, similar to roving effects in VPL.

However, an alternative interpretation to the stochasticity hypothesis is that the task difficulty was not matched between the RI and FI groups since the length of comparison intervals were selected from a normal distribution in the RI group, but a uniform distribution in the FI group. This entails that the majority of the comparison values in the RI group were very close to 200ms (194 and 206ms), thereby making discrimination more difficult than the FI group, who received a wide range of comparison values. Since learning is impaired in more difficult discrimination conditions (Ahissar & Hochstein, 1997), the selective improvement in performance for the FI group observed in Experiment 1 can be attributed to greater task difficulty for the RI group, rather than stochasticity in the stimulus structure (random vs. fixed).

To address this possibility, we conducted a second experiment with nine new participants. All stimuli and parameters of the task are identical to the RI condition in Experiment 1 with the exception that comparison values are now selected at random from a uniform distribution (with the same bounds as Experiment 1). With this manipulation, there is now an equal likelihood of being presented with a value close to 200ms than any other value between the minimum and maximum bounds. Therefore, Experiment 2 serves to equate the frequency of occurrence of all comparison intervals between the two groups. If our hypothesis is correct, we would expect to replicate our results of the RI group from Experiment 1.

## Methods (Experiment 2)

### Participants and Stimulus

Nine additional young adults were recruited from the same subject pool as Experiment 1 (8 females; mean age: 22.4 ± 2.9 years), none of which participated in any previous TPL experiments. One participant was excluded due to abnormally high thresholds (>2.5SD from the group mean), leaving eight participants for analyses. All stimulus parameters are identical to the RI condition of Experiment 1, with the exception that on every trial, a comparison interval was selected at random, with equal likelihood, from 158 to 194ms or from 206 to 242ms. The mean and standard deviation parameters for the FI (Exp. 1) and RI (Exp. 2) groups are identical. Consent was obtained prior to participation and approved by the IRB at Brown University.

## Results

Similar to Experiment 1, we fit individual psychometric functions for each subject for each day and calculated a change in DL between the first and last day of training. TPL as indexed by an improvement in discrimination performance for Experiment 2 was more similar to the RI group in Experiment than the FI group (Figure 3). A mixed-model ANOVA on the DLs with Experiment (3) and Session (2) as between-subjects factors only revealed a significant main effect of training, F_1,29_ = 5.52, p = .026, η^2^ = .03, but not experiment or interaction between experiment and training. Separate post-hoc pairwise t-tests were conducted for the FI and RI groups: there is a significant effect of training for the FI group, t_11_ = 2.77, p = .018, but not for the RI group, t_7_ = 0.59, p = .57. The lack of a significant interaction could be due to the smaller sample size (i.e., greater variability) in Experiment 2. Also consistent with our findings of Experiment 1, we did not observe a transfer of learning from the trained task to the untrained task (Figure 2). An omnibus ANOVA with Greenhouse-Geiser corrections again revealed a significant main effect of condition, F_3,87_ = 4.01, p = .038, η^2^ = .05 but no group by session by condition interaction, F_6,87_ = 1.39, p = .22 (all other p-values > .05).

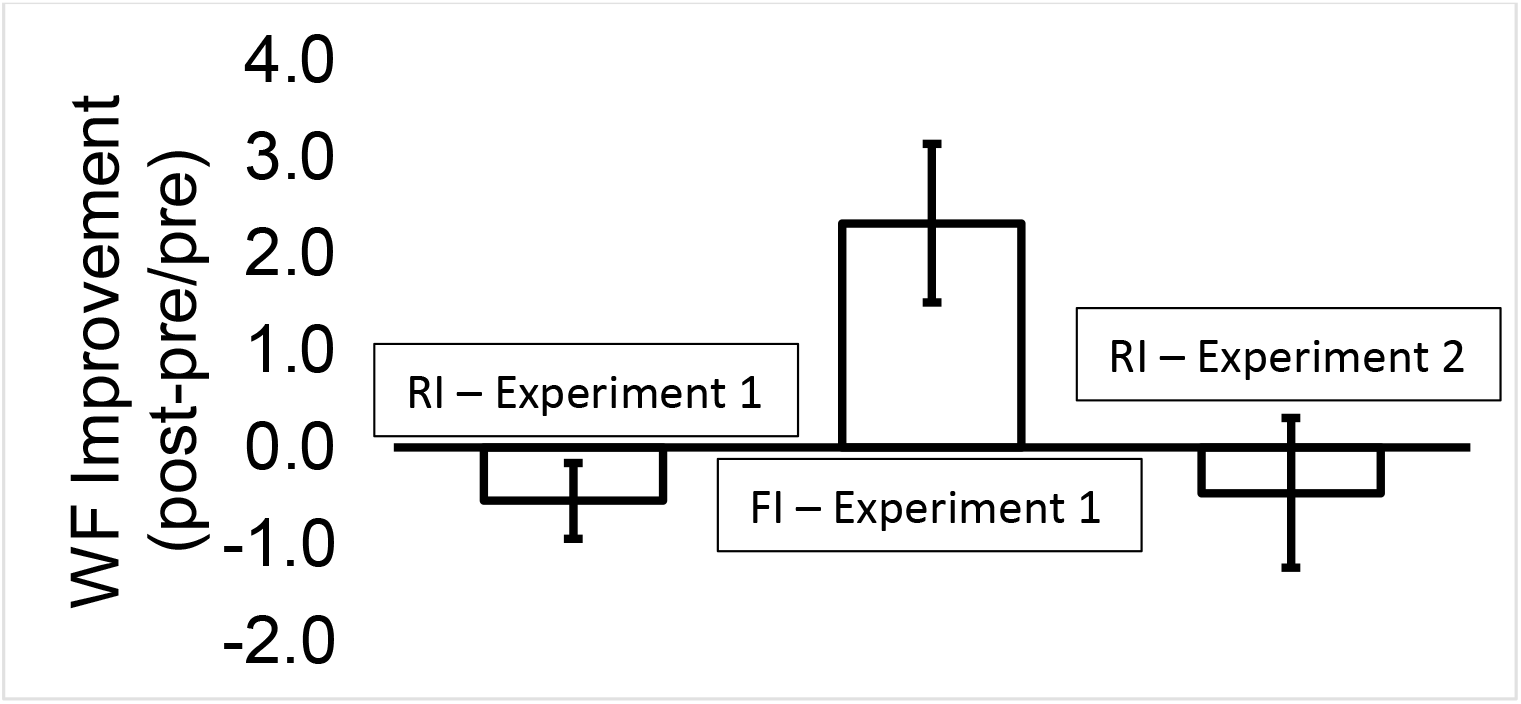

## General Discussion

A central question of TPL is what is actually being changed when we learn to time. Evidence of within-modality generalizations in TPL (e.g., Wright et al., 1997; Karmarkar & Buonomano, 2003) suggest that training improves our representation for a temporal interval (e.g., 100ms) which, after a more extended period of practice (Wright, Wilson, & Sabin, 2010), can improve discrimination on untrained dimensions of that interval (e.g., 100ms at a different pitch). In contrast to a representational hypothesis, results from our experiments suggest that TPL must also rely on changes in rule-based processing strategies which optimizes performance only in a task-relevant dimension. Whether TPL improves the representation of a temporal interval (low-level) or decision processes (high-level) or both, is the “locus of learning” problem of TPL. The goal of the present set of experiments was to provide evidence in support high-level changes in temporal learning.

In Experiment 1, we found that TPL of a single-interval comparison task was contingent on the input structure of information during training. Specifically, learning was impaired when random intervals were presented from trial to trial, similar to roving effects observed in VPL. On the other hand when a prefixed set of comparison intervals were used, a training-related reduction in discrimination threshold was observed between the pretest and posttest sessions. This finding stands in contrast to the low-level change hypothesis because the length of the standard interval (i.e., 200ms) was identical in all groups, therefore if the optimal strategy was to make a mental comparison between one’s internal representation of a 200ms interval and the presented comparison interval, learning should occur irrespective of whether *C* is random or fixed. The observation that only the FI group showed significant improvement in performance therefore suggests that individuals were not actually improving their representation of the 200ms interval but rather, they are learning the optimal decision strategy for the task, possibly strengthen the relevant stimulus-response relationships. For the FI group, this strategy was viable because there are only 8 possible comparison values, so only 4 stimulus-response pairs needed to be optimized. In contrast for the RI group, it was impossible to use the same strategy because there were an infinite number of possible values for *C*, with no repetition. Therefore, an alternative – albeit less successful – strategy must be adopted in the RI group. What our results suggest therefore, is that TPL does not rely exclusively on changes in low-level representations of an interval (Karmarkar & Buonomano, 2003; Nagarajan et al., 1998; van Wassenhove & Nagarajan, 2007); that top-down processes are also necessary in improving performance in a temporal discrimination task and it is this set of strategies that are improved with training rather than low-level changes in the representation of the trained (200ms) interval itself.

Further evidence in support of high-level changes in TPL is our second finding that learning on one temporal discrimination task is specific to the trained task structure and did not generalize to an untrained task using the same (i.e., learned) interval. In both experiments, we failed to find a generalization of learning from the trained task (single-interval comparison) to the untrained task (two-interval discrimination). This result suggests that learning may be bound by the greater attentional context of task-specific processes, manifested as an inability to generalize the high-level structure and decision rules learned in a task to a new, untrained task (which would necessitate a different set of response strategies). Taken together, our second finding of task-specificity is also consistent with high-level changes associated with TPL.

A number of past studies have observed a symmetric (Warm, Stutz, & Vassolo, 1975) transfer of TPL between the visual and auditory modalities, as well as between auditory and somatosensory/motor timing tasks (Meegan, Aslin, & Jacobs, 2000; Nagarajan et al., 1998). These generalizations are often taken to be evidence in support of the existence of a centralized timing mechanism which is improved with practice (see Ivry, Schlerf, & Ivry, 2008). While our finding of task-specificity across our two auditory discrimination tasks seem contradictory to the above findings, a number of studies have also found either limited (Bratzke, Seifried, & Ulrich, 2012) or no transfer of learning to untrained modalities (Grondin & Ulrich, 2011). Thus, it is unclear what mechanisms actually underlie the ability for learning to generalize across modalities, and in this case, across perceptual tasks.

One possible explanation for the task-specificity observed in our experiments is an asymmetric generalization of TPL between an easier and harder perceptual task. In a seminal study by Ahissar & Hochstein (1997), initial task difficulty during training plays a major role in determining the stimulus characteristics that are later generalized. This finding is later formalized as RHT (Ahissar et al., 2009; Ahissar & Hochstein, 2004; Hochstein & Ahissar, 2002) where perceptual learning is seen as an improvement in the *perception* rather than the *sensation* of a perceptual stimulus. As such, learning proceeds from easier tasks and gradually to harder tasks. However, training on a harder task induces a level of specificity that discourages the generalization of learning to an easier task (Ahissar & Hochstein, 1997). In the present study, it can be argued that the single-interval discrimination task has a greater cognitive involvement (i.e., comparing a single interval to a mental reference) in comparison to the two-interval discrimination task (i.e., two intervals in succession). If this is the case, we would expect a unidirectional transfer from the easier (two-interval discrimination) to harder (single-interval discrimination) task, but not vice versa, consistent with RHT.

Taken together, our two findings support the involvement of high-level changes in TPL. In addition, we demonstrated similarities between the perceptual learning of visual features and temporal intervals through roving and task-specificity. These findings are consistent with the idea that perceptual learning proceeds in a top-down manner, where task-relevant information is learned and only conditions which share temporal and task-specific characteristics become generalized. This is the first demonstration of task-specific learning in time perception and suggest that TPL at least partially reflects higher-level reweighting or decision unit changes. Our findings provide grounds for constructing a unifying framework of perceptual learning and address the dichotomy between sensory-stage and decisional-stage learning at a systematic level.

## Acknowledgments

This research was made possible by support from the Edgar J. Marston grant awarded to RMC (EN464266) and an R01EY027841 awarded to TW from the NIH. We would like to thank Daniela Ravasio for her help with data collection and David Freestone for helpful feedback on earlier drafts of this paper.

